# Obtaining 3D Super-resolution Information from 2D Super-resolution Images through a 2D-to-3D Transformation Algorithm

**DOI:** 10.1101/188060

**Authors:** Andrew Ruba, Wangxi Luo, Joseph Kelich, Weidong Yang

## Abstract

Currently, it is highly desirable but still challenging to obtain three-dimensional (3D) superresolution information of structures in fixed specimens as well as dynamic processes in live cells with a high spatiotemporal resolution. Here we introduce an approach, without using 3D superresolution microscopy or real-time 3D particle tracking, to achieve 3D sub-diffraction-limited information with a spatial resolution of ≤ 1 nm. This is a post-localization analysis that transforms 2D super-resolution images or 2D single-molecule localization distributions into their corresponding 3D spatial probability information. The method has been successfully applied to obtain structural and functional information for 25-300 nm sub-cellular organelles that have rotational symmetry. In this article, we will provide a comprehensive analysis of this method by using experimental data and computational simulations.

## Introduction

Since stated by Ernst Abbe in 1873, the resolution of conventional light microscopy has been limited to approximately 200 nanometers laterally (x,y) and 600 nanometers axially (z) due to light diffraction from the microscope objective^1,2^. Currently, super-resolution light microscopy techniques break this limitation and allow for the capture of static or dynamic images with subdiffraction resolution (< 200 nm) in all three axes. The techniques generally fall into two broad categories: optical based approaches such as stimulated emission depletion (STED) microscopy, which generate a sub-diffraction illumination volume due to the nonlinear optical response of fluorophores in samples through laser modifications; and single-molecule based mathematical approaches such as photoactivated light microscopy (PALM) and stochastic optical reconstruction microscopy (STORM). PALM and STROM utilize mathematical functions to localize the centroids of fluorophores and then reconstitute these centroids to form super-resolution images^1–4^. Although these super-resolution techniques have revolutionized imaging of biological samples via unprecedented spatial resolution, they are still limited in acquisition time (seconds to hours) and axial spatial resolution (typically > 50-100 nm)^1–4^. Meanwhile, fast, three-dimensional (3D) superresolution imaging is critical for obtaining structural or dynamic information in live cells, which are inherently 3D objects. Moreover, many biological functions in sub-cellular organelles are near or below the spatiotemporal resolution limit of current 3D super-resolution imaging techniques, such as nucleocytoplasmic transport through 50-nm nuclear pore channels with millisecond transport times^5–7^.

Typically, 3D super-resolution imaging is more technically demanding than 2D super-resolution imaging. This is due to the fact that the point spread function (PSF) of the emitting fluorescent probe in the axial dimension is much larger than in the lateral dimension at the focal plane of the light microscopy objective^1^. Several methods have been developed to improve axial resolution in fluorescence microscopy. One category is to alter the shape of the PSF of the fluorescent probe along the optical axial position and then determine the probe’s axial information by referencing a pre-determined relationship (setup through control experiments) between the shape of the PSF and the corresponding locations in the z dimension^8^. The other is to use two objectives to improve the axial resolution after comparing fluorescent signals of the probes from these objectives with or without interference^9^. Typically the above approaches involve expensive and complex optical implements. Here we introduce an alternative approach for overcoming this axial dimension resolution limitation that does not require unique optics. Instead of modifying the microscopy setup, this approach is a post-localization analysis that transforms 2D super-resolution images or 2D single-molecule localization distributions into their corresponding computational 3D probability information. This is done by utilizing biological structures that are inherently radially symmetrical in one of the three dimensions. The whole process includes three major steps: 1) determine the structure of a sub-cellular organelle by electron or expansion microscopy^10,11^; 2) conduct single-particle tracking and/or single-molecule localization to get 2D super-resolution images or 2D single-molecule localization distributions in the structure^12,13^; and 3) transform these experimentally obtained 2D data into their corresponding computational 3D probability information by utilizing the previously determined rotationally symmetrical structure. Here, we will provide a detailed analysis of the 2D-to-3D transformation process and demonstrate its applications in determining 3D structural and functional information in sub-cellular submicrometer organelles that have rotational symmetry^14,15^, such as the nuclear pore complex (NPC) and the primary cilia^5,6,16–18^. Also, we will further demonstrate that the 2D-to-3D transformation process can be extended to convert 2D super-resolution images obtained from currently existing 2D super-resolution light microscopy techniques, by using STORM-based 2D data of microtubules as an example.

## Results

### Mathematical concept and detailed process for the 2D-to-3D transformation algorithm

As shown in Fig. 1 and Online Methods, the main idea behind the 2D-to-3D transformation algorithm is that, for any radially symmetric biological structure (typically determined by using electron microscopy) such as NPCs, primary cilium and microtubules seen in Figure 1A-D, an area matrix in the radial dimension can be developed (y and z dimensions in Figure 1E). As molecules locate in or travel through these structures, their locations can be projected into the xy or the yz plane, depending on whether microscopy imaging of the structure is conducted at the lateral or the axial dimension respectively (Fig. 1F). Typically, in practice it is much easier to obtain 2D superresolution images of these targeted molecules in the xy plane than the yz plane because of the much larger axial dimension than the radial dimension in these structures. In the end however, the 3D information on the structure can either be collected in the xy and yz planes. After further projecting the 2D molecular spatial locations into the y dimension from either the xy or the yz plane, the obtained two y-dimension histograms in principle are identical as demonstrated in Fig. 1G. Then, based on the two identical y-dimension histograms, each column in the y-dimension histogram projected from the xy plane will be equal to the areas times the densities for each radial bin of the yz plane (Fig. 1 G-H). Finally, as detailed in the mathematical calculation shown in Online Methods, the densities in the radial dimension can be obtained by solving the matrix equations, which eventually reconstitute the corresponding 3D super-resolution information for the structure (Fig. 1 I-K).

**Figure 1.**
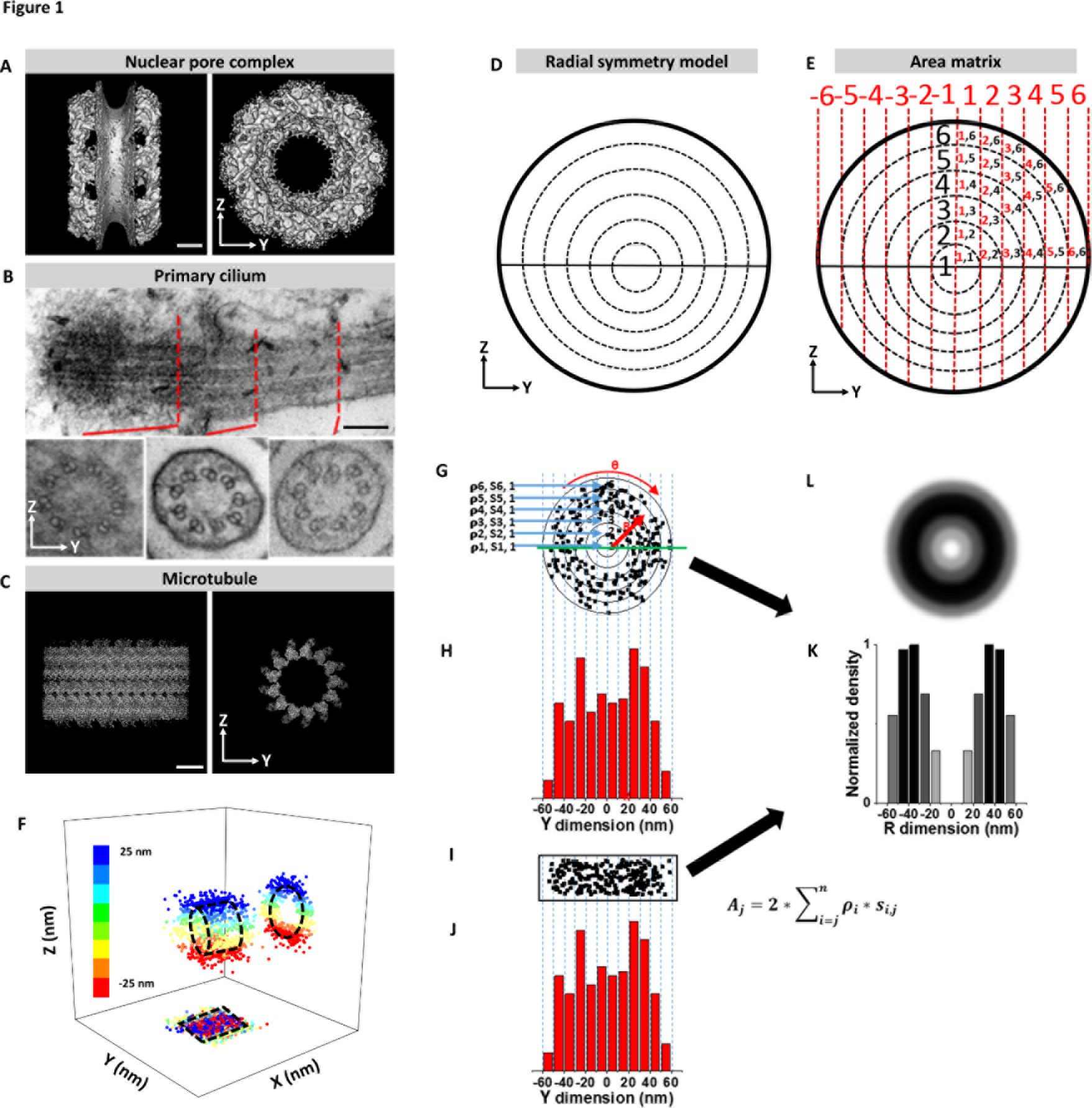
An area matrix can be developed for radially symmetric biological structures that reflects 2D single molecule data. **(A)** 3D electron tomography image of the nuclear pore complex averaged rotationally with 8-fold symmetry^46,47^. Scale bar = 20 nm. **(B)** Transverse slice of the primary cilia using transmission electron microscopy^47,48,49^. From left to right, the red lines indicate cross-sections of the basal body, transition zone, and ciliary shaft of the primary cilium. Scale bar = 100 nm. **(C)** 3D electron tomography image of a microtubule averaged rotationally with 13-fold symmetry^47,50,51^. Scale bar = 10 nm. **(D)** Structures from (A), (B), and (C) may be simplified as a radially symmetric circle in the y and z dimensions. **(E)** An area matrix may be designed where the radially symmetrical simplified model is divided along the y dimension due to the fact that the projection of 2D single molecule data is along the y dimension even though it came from the y and z dimensions. **(F)** Simulated single molecule data in the x, y, and z dimensions from a cylinder with ideal radius of 25 nm and localization error of 5 nm for demonstration of the 2D-to-3D density transformation. Dotted lines represent the ideal structure from which the simulated originated. **(G)** Single molecule data from (F) in the y and z dimensions superimposed with the area matrix from (E). **(H)** Histogram of the single molecule data from (G) along the y dimension to model the effects of 2D microscopy, from which a y dimensional histogram can also be obtained. **(I)** Single molecule data from (F) in the x and y dimensions. **(J)** Histogram of the single molecule data from (I) along the y dimension. **(K)** The resultant radial density map of the single molecule data from (G) determined by dividing the number of points in each radial bin by the area of each radial bin as well as the resultant radial density map of only the y dimensional histogram from (I) determined by using the 2D-to-3D density transformation. **(L)** Radial density map from (K) expressed as a cartoon reconstruction of the density of single molecules through the original 3D structure. The intensity of rings, from black to white, indicates highest to lowest normalized density.

### Monte Carlo simulation demonstrates the parameters required for accurate 3D density map reconstruction

Next, we use Monte Carlo simulation^19,20^ to demonstrate that two critical parameters, the single-molecule localization error of targeted molecules^21^ and the number of single-molecule locations^4,22^, will determine the reproducibility of obtaining accurate 3D superresolution information in the biological structures with rotational symmetry. Typically, the spatial localization of individual targeted molecules labeled with fluorophores or fluorescent proteins was determined with non-zero localization error because of, primarily, background noise and limited photon collection in real experiments^21^. To mimic typical transport routes of proteins in the NPC or primary cilia, our Monte Carlo simulations were performed where varying numbers of singlemolecule locations were randomly simulated on an ideal radius (R_I_) (Supplemental Figure 3A-B). Then, single-molecule localization error (σ_LE_) was added to R_I_ by sampling an error value from a normal distribution with a standard deviation of σ_LE_ (Supplemental Figure 3C). Subsequently, the 2D-to-3D transformation algorithm was performed on only the y dimensional data of the simulated single-molecule localization distribution around R_I_ to model the loss of z dimensional information during the 2D microscopy projection process. The peak position of the transformed 3D density histogram was then determined by Gaussian fitting to produce a measured mean radius (R_M_) which may deviate from the R_I_ due to limited number of simulated locations and non-zero singlemolecule localization error (Supplemental Figure 3D). We conducted 10,000 iterations of this process and obtained 10,000 R_M_ values, in which the mean of the R_M_ values converges on R_I_ as expected (Supplemental Figure 3E). To quantify how reproducible a single experimental data set is, we set out to determine how many individual R_M_ values from the whole distribution of R_M_ fell within an acceptable range of R_I_. The acceptable range was defined as the R_I_ ± σ_LE_ because, in principle, any single R_M_ value can only be accurately localized within the range of approximately two standard deviations of its Gaussian fitting, similar to the concept of resolution stated by the Rayleigh Criterion^23^ (Supplemental Figure 3F). We expect that a high number of simulated singlemolecule locations or low single-molecule localization errors would increase the number of iterations that fall within the acceptable range, thus resulting in a high reproducibility rate. To determine the reproducibility rate, we found that two critical steps should be correctly followed first in the process: first, optimize the bin size for each set of simulation parameters. This is accomplished by determining the smallest bin size that produces no statistical difference by Chi square analysis between the original 2D histogram and the back-calculated 2D histogram obtained by multiplying the 3D density histogram by the corresponding area matrix (Supplemental Figure 4). The second step is to account for the sensitivity of the inner bins of the area matrix when determining the accurate Rm peak fitting (Supplemental Figure 5).

First, to test the effects of single-molecule localization error on the final R dimensional peak fitting obtained for the 3D transformed density histogram, Monte Carlo simulations were performed with an R_I_ of 25 nm, data point number of 1,000,000, and σ_LE_ ranging from 0 to 30 nm (Figure 2A-D). In principle, as the single-molecule localization error becomes excessively large, the peaks in the 3D density map will become heavily overlapped on the y dimensional axis and, subsequently, the radial axis after the transformation algorithm. This will obscure the peak at R_I_ and make it indistinguishable. A ratio between the error of a bimodal Gaussian fitting and the error of a single Gaussian fitting is used to determine the indistinguishable overlap (Figure 2E-H). As shown in Figure 2I, a series of tests indicate that the bimodal fitting error becomes much larger than the single peak fitting error beyond a 21-nm localization error. This suggests that the experimental localization error cannot exceed 21 nm for any structure containing a transport route with a radius of 25 nm. While, this is smaller than the theoretical single-molecule localization error of 25 nm predicted by the Rayleigh Criterion, mainly due to the aforementioned sensitivity of the inner bins of the area matrix in this 2D-to-3D transformation process (Supplemental Figure 5). Moreover, the above results can be generalized by using a radius/precision (R/P) ratio to estimate whether the transport route can be distinguished before computational simulations or real experiments. As shown in Figure 2 C,G, the threshold case of a R/P ratio of 1.19 (25 nm/21 nm) presents the minimally distinguishable transport route in the R dimension, corresponding to a separation of ~68% of the single-molecule density around the radius. As mentioned above, 1.19 represents the case where the radius is approximately equal to the precision or standard deviation of the transport route; thus, separation of the radial distribution by ~1 standard deviation results in ~68% of single-molecules density separation. Meanwhile, when the R/P ratio is ≥ 2.0, a much higher degree (correspondingly ≥ 95% of single-molecule density) of peak separation and a well distinguished transport route can be achieved (Fig. 2 B, F).

**Figure 2.**
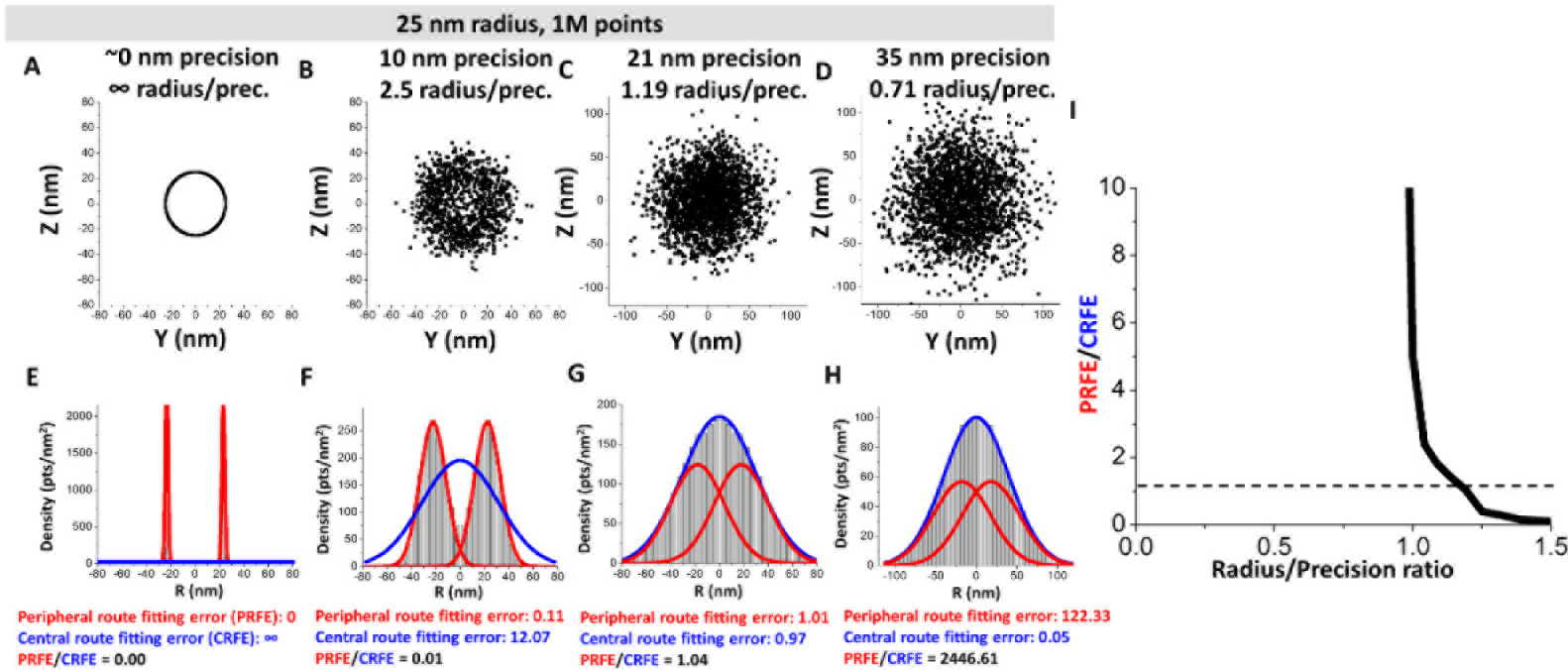
Varying the simulated precision shows the resolution limit of the 3D transformation algorithm. **(A), (B), (C), and (D)** 1,000,000 locations were simulated on an ideal radius (R_I_) of 25 nm with localization errors (σ_LE_) of 0, 10, 21, and 35 nm respectively. Scatter plots were downsampled to 2,000 locations for visualization. **(E), (F), (G), and (H)** The corresponding 3D transformed density histogram of each simulation from (A), (C), (E), and (G) respectively. To determine whether the density peaks could be distinguished after a given localization error had been added to the ideal data, the ratio between the fitting error of a bimodal Gaussian distribution (PRFE) and a single Gaussian distribution was used (CRFE). If the bimodal fitting error was less, then the peaks in the 3D transformed distribution can likely be distinguished. If the single peak fitting error was less, the localization error was too great and the two peaks were indistinguishable. **(I)** Up to 21 nm, the PRFE was much less than the CRFE. Above 21 nm, the CRFE was much less than the PRFE. Therefore, the maximum localization error allowed to distinguish a 25 nm ideal radius is ~21 nm. The results are generalized as precision/radius.

Next, using the R/P ratio of ≥ 2.0, we sought to determine the effects of the quantity of single molecule locations on constituting an accurate 3D transformed structure. To accomplish this, Monte Carlo simulations were performed with an R_I_ of 25 nm, σ_LE_ of 10 nm (the R/P ratio of 2.5), and data point numbers ranging from 50 to 1,000. Representative simulations are shown for 50, 100, 500 and 1,000 points, and the corresponding reproducibility percentages were calculated for each point number after 10,000 iterations (Figure 3A-H). Remarkably, only 100 and 350 points are sufficient to achieve 90% and 99% reproducibility respectively (Figure 3I). It is highly feasible to obtain 100-1,000 points experimentally, although the number of points is higher than the minimum Nyquist Sampling theorem estimation of 38 single-molecule locations^22^ (Supplementary Information) because of a non-uniform distribution of locations through the area matrix. Of note, we found that 1,000 single-molecule locations can already result in an accuracy of 1 nm in obtaining 3D transformed density map (Figure 3D, H, I). Thus in theory, given enough singlemolecule data, the accuracy could be unboundedly small as long as the R/P ratio is above 2. Finally, in some cases, we noticed that, if the width of transport route (2σ_w_, the width at one standard deviation of its Gaussian distribution) is significantly bigger than the single-molecule localization precision, the above R/P ratio should be modified as R/P_w_ (*P*_*W*_= *σ*_*w*_) before determining the final radius of the transport route and the number of single-molecule locations needed for a high reproducibility rate. This discrepancy between the single molecule localization precision and *σ*_*w*_ is due to a non-negligible width of the biological transport route.

**Figure 3.**
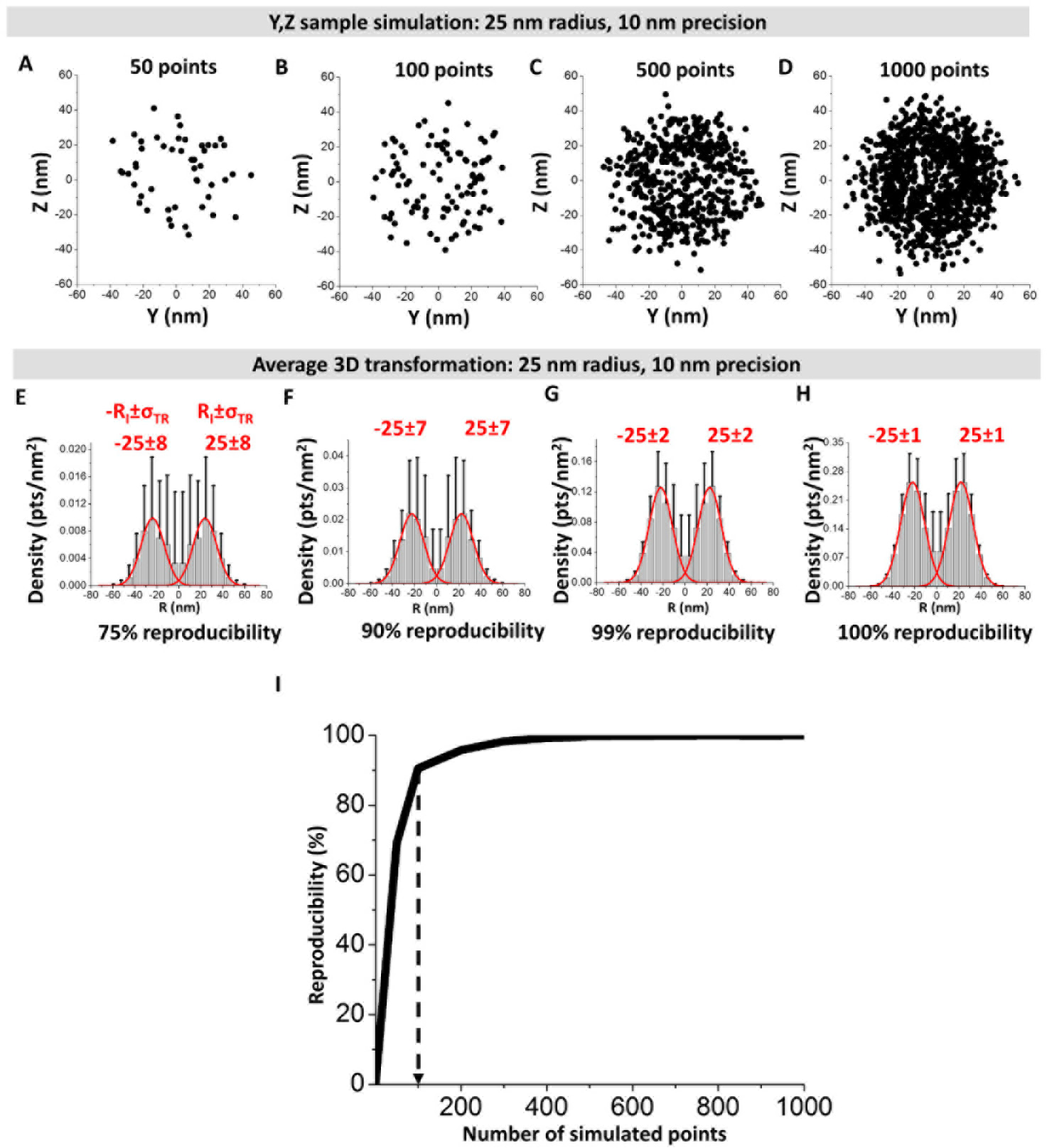
Varying the number of simulated points shows the effect of sampling error on the peak fitting of the 3D transformed data. **(A), (B), (C), and (D)** 50, 100, 500, and 1,000 locations were simulated on an ideal radius (R_I_) of 25 nm with localization error (σ_LE_) of 10 nm. **(E), (F), (G), and (H)** The average 3D transformation of 10,000 simulated data sets for each number of simulated locations. Error bars indicate standard deviation of each histogram bin value while Ri±otr indicates the average peak fitting ± the standard deviation of the peak fittings. That is, the standard deviation (σ_TR_) of the R_M_ distribution outlined in Figure 1E. The reproducibility percentage, the number of peak fittings that fell within the localization error, is shown beneath each 3D transformed density histogram. **(I)** The plot of reproducibility percentage for 10,000 iterations of each number of points up to 1,000. The simulation parameters were the same as (A),(B), (C), and (D) except for the varying point number.

**Figure 4.**
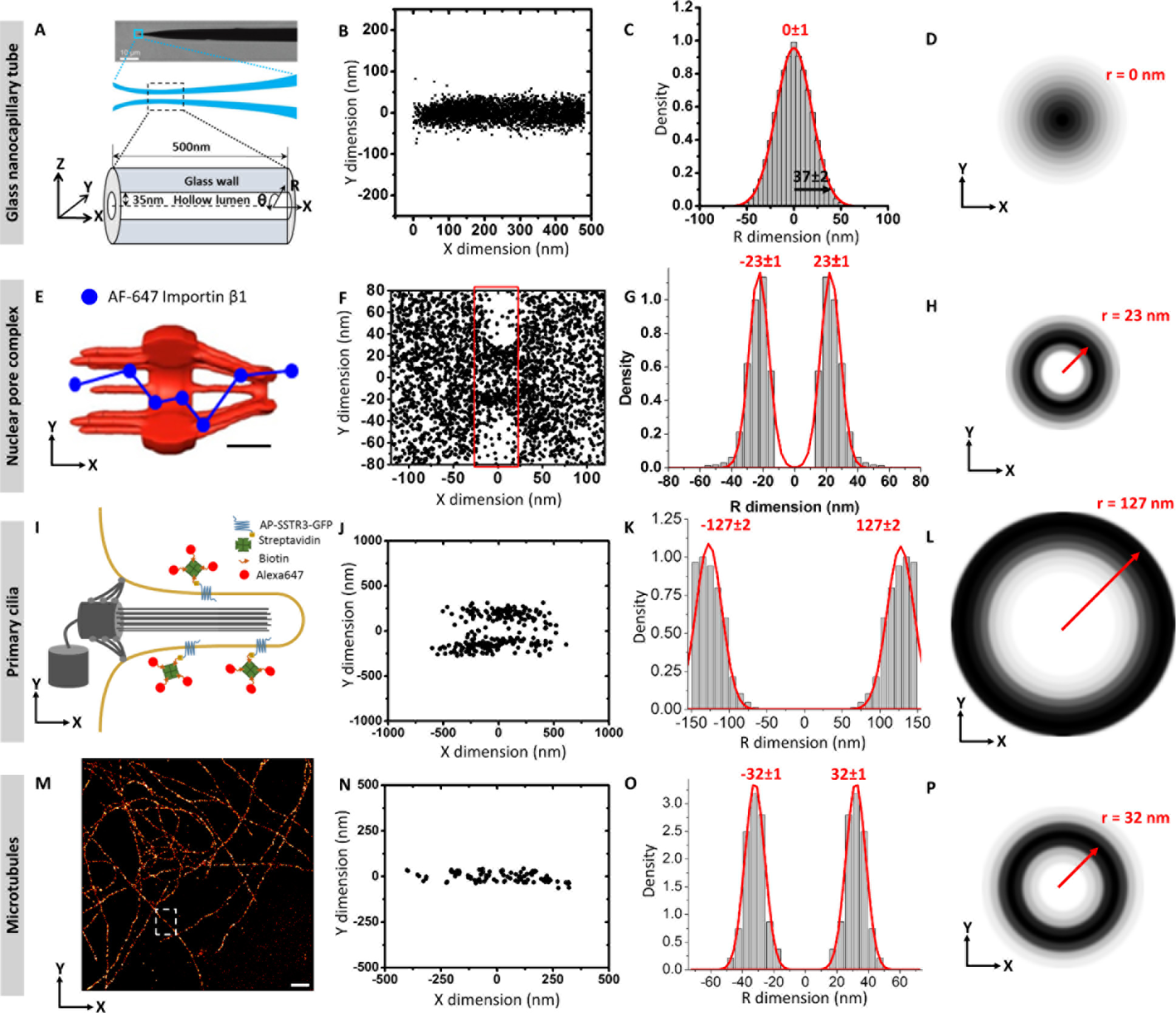
3D density maps accurately reconstruct the locations of labeled molecules in various structures. **(A)** Schematic of the glass nanocapillary tube with dimensions of 35 nm inner radius. **(B)** x and y dimensional single molecule data from the tracking of Alexa Fluor 647 inside the glass nanocapillary tube. **(C)** 3D density map of (B) showing the width at 2 standard deviations from the mean is ~ 37 nm. **(D)** Radial density map from (C). **(E)** Schematic of the nuclear pore complex with a model trajectory for Alexa Fluor 647-labeled Importin β1. Scale bar = 50 nm. **(F)** x and y dimensional single molecule data from the tracking of Alexa Fluor 647-labeled Importin β1 in the nuclear pore complex. **(G)** 3D density map of (F) showing the peak fitting of the Importin β1 transport route is ~ 23 nm. **(H)** Radial density map from (G). **(I)** Schematic showing the Alexa Fluor 647 externally-labeled SSTR3 in the shaft of a primary cilium. **(J)** x and y dimensional single molecule data from the tracking of externally-labeled SSTR3 in the primary cilium. **(K)** 3D density map of (J) showing the peak fitting of the SSTR3 transport route is ~ 127 nm. **(L)** Radial density map from (K). **(M)** 2D STORM data showing the reconstructed super-resolution image of tubulin labeled with a primary and secondary antibody attached to Alexa Fluor 647. Scale bar = 1 µm. **(N)** x and y dimensional single molecule data from section of 2D STORM data in the dashed white box from (M). **(O)** 3D density map of (K) showing the peak fitting of the labeled tubulin from (N) to be ~ 32 nm. **(P)** Radial density map from (O).

### SPEED Microscopy

Our lab has previously developed SPEED microscopy to fill the technique niche of capturing single molecules transporting through sub-diffraction-limit bio-channels at high spatial (< 10 nm) and temporal (< 1ms) resolution^14,15^. We achieve this through four main technical modifications on traditional epifluorescence or confocal light microscopy. (1) A small inclined or vertical illumination PSF is used for the excitation of single transiting molecules through bio-channels in the focal plane. This greatly increases the allowable detection speed (up to 0.2 ms per frame for the CCD camera we used) by reducing the number of camera pixels required for detection. Also, it significantly avoids out-of-focus fluorescence with an inclined illumination PSF in a similar way as total internal reflection microscopy^24^. (2) A high optical density (100-500 kW/cm^2^) in the small illumination volume causes a high number of photons from the fluorophores to be emitted. (3) A collection of a high-number of photons in a short time period increases spatial resolution by greatly reducing the effects of molecular diffusion on the single-molecule localization precision of moving molecules^21,25^ (Online Methods). (4) Pinpointed illumination in live cells causes negligible photo-induced toxicity^26–28^. Thus, SPEED microscopy meets the needs of high 2D spatiotemporal resolution for *in vivo* single-molecule tracking in dynamic bio-channels such as single NPCs. However, SPEED does not directly obtain any 3D information. As demonstrated below, we have employed SPEED microcopy to obtain 2D single-molecule data in glass nanocapillary tubes, the NPC, and primary cilia. However, other single-particle tracking or super-resolution microscopy approaches may be employed to obtain similar 2D data sets for processing by the 2D-to-3D transformation.

### Experimental validation of 2D-to-3D transformation process in several systems: glass nanocapillary tube, nuclear pore complex, primary cilia, and microtubules

Since the 2D-to-3D transformation algorithm requires radial symmetry, we first used an ideal artificial glass nanocapillary to test the algorithm’s accuracy when coupled with SPEED microscopy for data acquisition. The glass nanocapillaries (GNCs) were fabricated using laser-assisted capillary-pulling of quartz micropipettes which can generate pore diameters ranging from 20 nm to 300 nm. The dimensions of GNCs used in this study were determined by Helium scanning transmission ion microscopy to have an inner radius ~35 nm^29,30^. With that parameter in mind, the dimensions of the GNC were re-measured by determining the 3D density map of 1-nM Alexa Fluor 647 that was pumped into the inner lumen of the GNC. After thousands of 2D spatial locations for Alexa Fluor 647 were collected with a single-molecule localization precision of ≤ 5 nm, the final 3D density map revealed a radius of 37±2 nm (a width at the two standard deviations of the Gaussian function), agreeing well with the 35-nm inner radius of the GNC imaged by helium ion microscopy with a reproducibility rate of 100% and a route localization error of 0.25 nm (Fig. 4 A-C).

After the inner diameter of the GNC was confirmed by the application of the 2D-to-3D transformation algorithm, we moved to two different macromolecular trafficking in sub-cellular organelles: Importin β1 (a major transport receptor in nucleocytoplasmic transport^31–33^) moving through the NPC and externally-labeled SSTR3 (a major transmembrane protein in primary cilia^34–36^) on the surface of primary cilia on the ciliary shaft. Previously, our lab has revealed that Importin β1 assists the movement of protein cargo via interactions at the periphery of the NPC, a selective gate between the cytoplasm and nucleus^14,15^. In this analysis, we present a total of 450 2D spatial locations with a single-molecule localization precision of < 10 nm at the NPC's scaffold region for Importin β1 within the NPC. The corresponding 3D density map clearly shows a high density region for Importin β1 at 23±1 nm along the radius of the NPC with a 100% reproducibility rate and a route localization error of ~1 nm (Fig. 4 D-F). Similarly in primary cilia, SSTR3, was externally-labeled with Alexa Fluor 647^37^ and tracked using SPEED microscopy along the length of primary cilia, ~125 nm in radius determined by electron microscopy (EM)^^16–18^^. Agreeing well with the EM-determined diameter, 260 externally-labeled SSTR3 molecules with a singlemolecule localization precision of ≤ 10 nm were determined to have a high density region at 127±2 nm along the radius of primary cilia, with a 97% reproducibility rate and a route localization error of ~2 nm (Fig. 4 G-I). It is noteworthy that the ~35-nm width of SSTR3’s route that is much bigger than the single-molecule localization precision resulted in the lower reproducibility rate and the route localization error when compared to the determinations in the GNC and NPC.

Lastly, to test whether the 2D-to-3D density transformation algorithm could be applied beyond SPEED microscopy, we measured the diameter of microtubules^38–40^, in which tubulins were labeled by a primary and secondary antibody conjugated to Alexa Fluor 647 and then imaged by 2D-STORM^41^. By converting the published 2D super-resolution image for a microtubule (112 single-molecule locations) into its 3D structure by our 2D-to-3D transformation algorithm, we determined the diameter of microtubules in this specific sample to be 64±1 nm with a reproducibility rate of 90% and a route localization error of ~1 nm (Fig. 4 J-L). This result agrees well with previous determinations by using EM^42^ and 3D super-resolution microscopy^1^.

## Discussion

In this paper, we presented a detailed analysis of the 2D-to-3D transformation algorithm that enabled us to obtain 3D super-resolution information from 2D super-resolution images or 2D single-molecule localization data without using 3D light microscopy setups. The roles that two critical factors played in reproducing the accurate 3D super-resolution information, the single molecule localization error and the number of single molecule locations, have been fully discussed. We also discussed a general rule that a transport route can be well distinguished if it has a radius/precision ratio greater than 2 and the ratio-based minimum number of single-molecule locations (Supplementary Figure 6). The successful applications in various systems, including the GNC, the NPC and primary cilia in live cells, and microtubules in fixed samples, prove the robustness of achieving accurate 3D super-resolution information by combining 2D experimental data and the 2D-to-3D transformation algorithm.

It is noteworthy that one prerequisite of the algorithm is that the density of the molecules of interest is constant along a given radial bin for the biological structure. Normally, transmission electron microscopy is appropriate for determining the structure and its rotational symmetry with high spatial resolution. Another alternative approach is expansion microscopy, by which the size of a specimen can be enlarged 4.5 to 20 times without significant distortion and the enlarged structure can be labeled and imaged by epi-fluorescence or confocal light microscopy. As demonstrated, with the known structures of the NPC, primary cilia, and microtubules, area matrices have been developed for the cylindrical structures based on their rotational symmetry for 2D-to-3D transformation algorithm. Furthermore, area matrices could also be developed for other regular or even irregularly-shaped, radially symmetric structures, with the only requisition of a constant density of the molecules of interest along a given radial bin.

## Online Methods

### Calculating the regions of the area matrix

In the 2D-to-3D density transformation process, we use an area matrix to reflect the contribution of each ring to the 2D distribution. As shown below, we define i as the radial number and j as the axial number. The density of single molecule locations in the same radius i is supposed to be uniform given the radial symmetry and the density is labeled as *ρ*_*i*_ here. The cross-ection of radial number i and axial number j is defined as A_(i,j)_. In the following equations, Δ*r* is the bin size, *ρ*_*i*_ is the density of molecules in the *i*th ring, *h* is the half of illumination depth and *ρ* is the constant background density outside the region of interest.

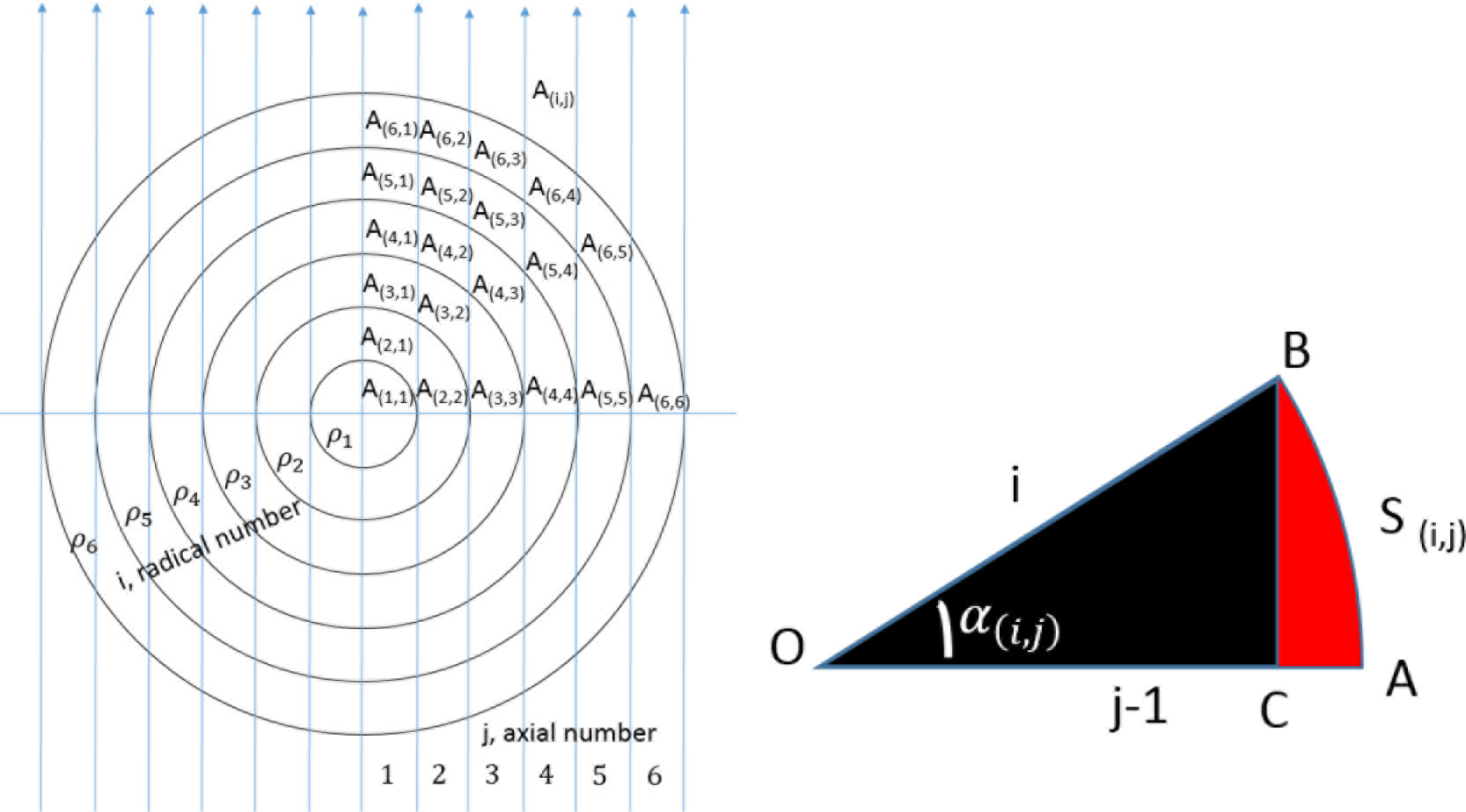

To determine the area of each sub-region (A_(i,j)_), we always start to calculate the area of the fan-shape area at *j*=*i*, in which:

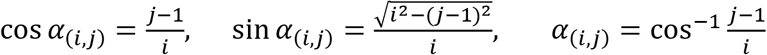

The red-shaded area is defined as S_(i,j)_:

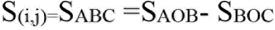

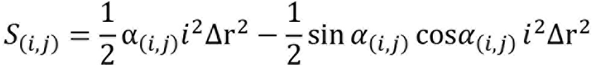

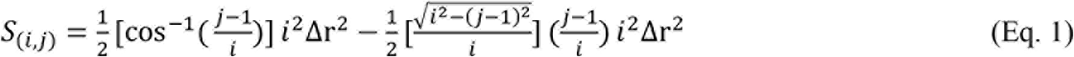

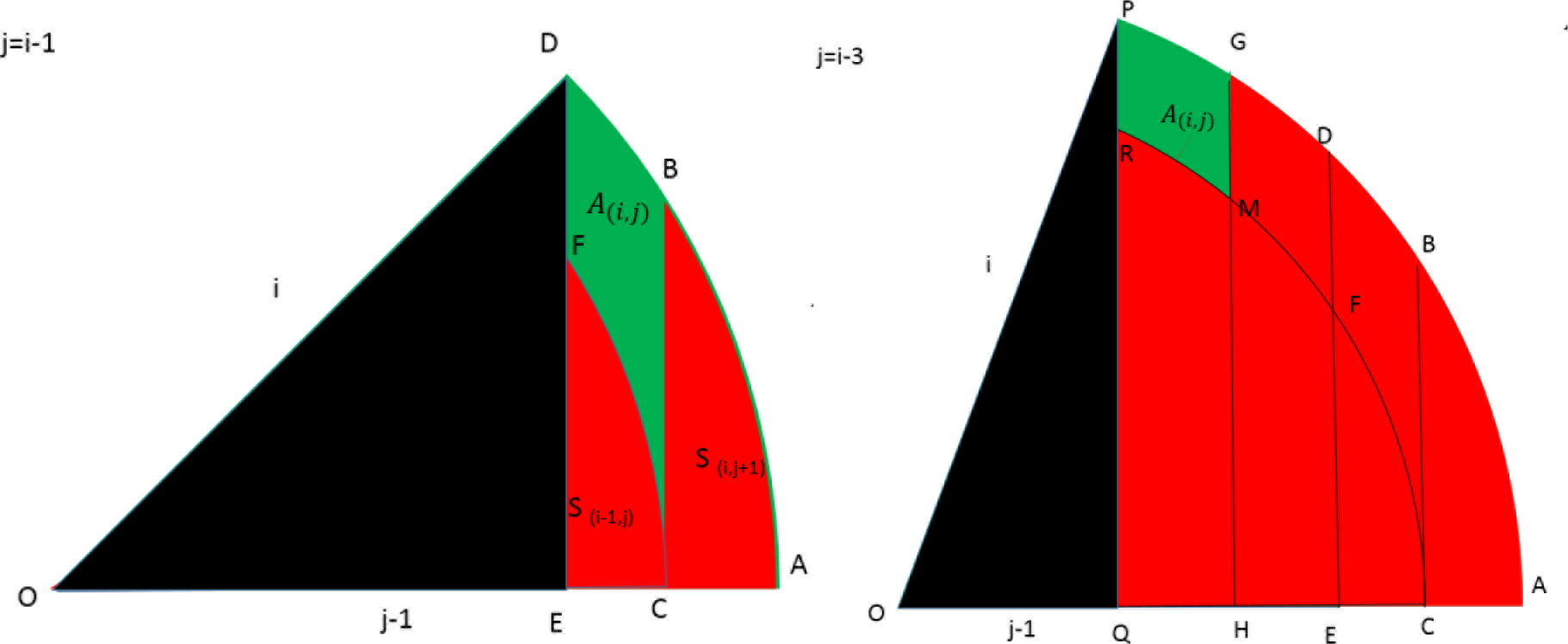

When *j* ≠ *i and i* > *j*, we need to calculate the area of the green-shaded region as follows:

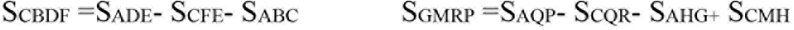

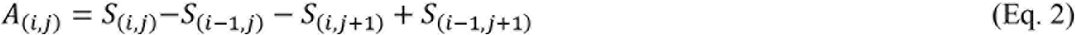

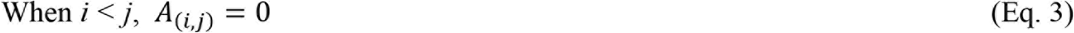

Following the above equations, all *A*_(*i*,*j*)_ can be precisely calculated.

Finally, N_j_, the number of events detected in the j^th^ column can be calculated with the following equation:

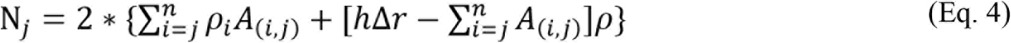

As soon as N_j_ and *A*_(*i*,*j*)_ are known, *ρ*_*i*_ will be calculated.

### Single molecule localization precision

For immobile molecules or fluorescent nuclear pores, the fluorescent spot was fitted to a 2D symmetrical or an elliptical Gaussian function, respectively, and the localization precision was determined by the standard deviation (s.d) of multiple measurements of the central point. So, the precision is presented as mean ± s.d. However, for moving molecules, the influence of particle motion during image acquisition should be considered in the determination of localization precision. In detail, the localization precision for moving substrates (σ) was determined by the formula 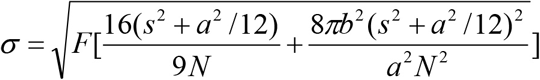, where F is equal to 2, N is the number of collected photons, a is the effective pixel size of the detector, b is the standard deviation of the background in photons per pixel, and 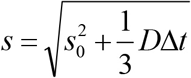, s_0_ is the standard deviation of the point spread function in the focal plane, D is the diffusion coefficient of substrate and Δt is the image acquisition time ^21,25,43–45^. In our experiments, more than 1,100 signal photons were collected from each targeted moving molecule. As a result, using the NPCs as an example, the localization precision for transiting molecules through the NPCs is calculated to be < 10 nm based on the above equations while the parameters were determined experimentally (N >1,100, a=240 nm, b≈2, s_0_=150±50 nm, Δ*t*=*0.4 ms*, D is < 0.1μm^2^/s for the tested substrates in the central channel of nuclear pores)^14,15^.

---

**Supplementary Figure 1.**
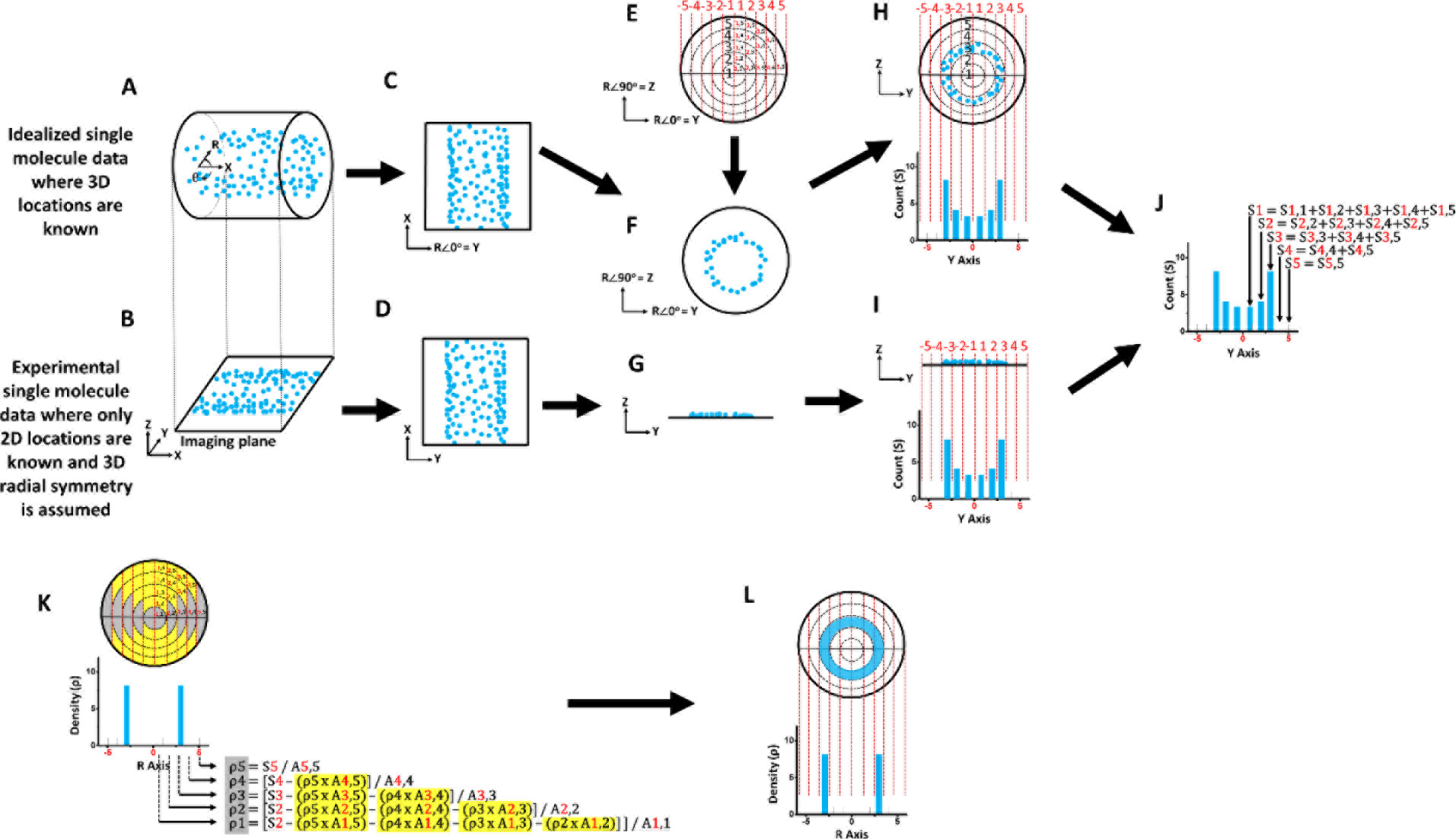
Further demonstration of 2D to 3D transformation process. **(A)** Idealized simulated single molecule data in x, y, and z dimensions. **(B)** Single molecule data from (A) projected to just the x and y dimensions, as it would be during the microscopy imaging process. **(C) and (D)** In the x and y dimensions, both the 3D single molecule data and the 2D projected data are identical. **(E)** An area matrix subdivided radially and axially to account for the radially symmetric structure from which the single molecule data originated. **(F)** Single molecule data from (A) viewed in the y and z dimensions. **(G)** Single molecule data from (B) viewed in the y and z dimensions. Since the data is projected, the z dimension is uniform. **(H) and (I)** y dimensional histograms from (F) and (G) are identical due to the fact that the z dimension is projected during the histogram process. **(J)** Each column in the y dimensional histogram in both (H) and (I) originate from the sum of the corresponding axial bins in the area matrix. **(K)** The 3D density histogram is calculated by using the outermost radial bin as a reference, calculating its density, then subtracting its contribution from the inner axial bins. The process is then repeated for the next most outer radial bin until the inner most radial bin is reached. **(L)** The final 3D density map gives a map of single molecule radial density rather than information about the 3D location of each single molecule.

**Supplementary Figure 2.**
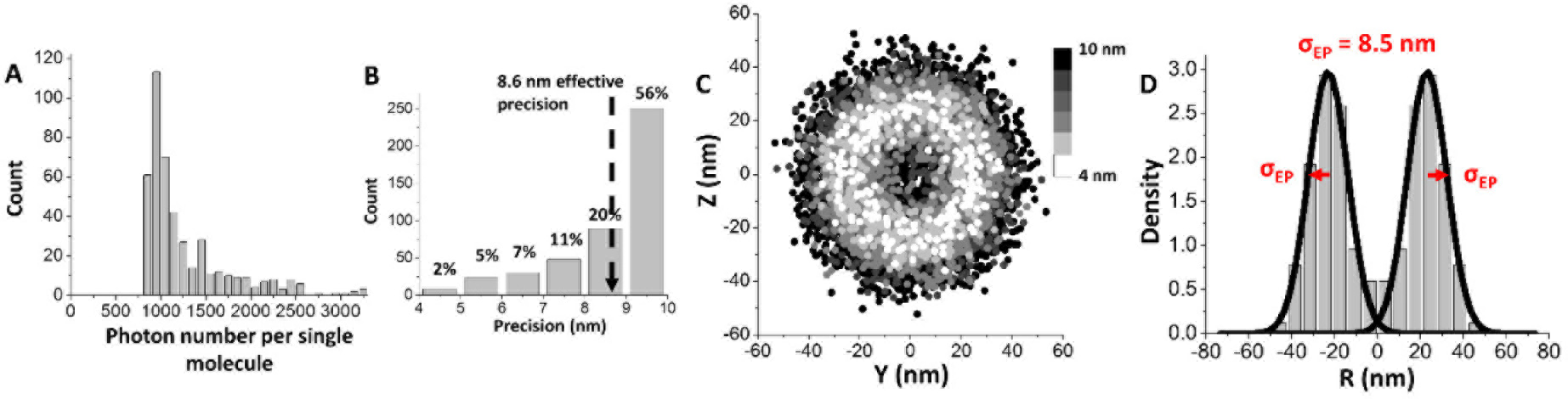
Minimum amount of experimental data points needed for a reliable 3D distribution of Imp β1 in the NPC. **(A)** During the data selection process, single molecules with less than ~800-1000 photons are removed before the final 3D transformation analysis. **(B)** In the precision distribution, this corresponds to a cutoff precision of 10 nm. Each whole number precision value below 10 nm threshold contributes a certain percentage to the overall weighted average precision of 8.6 nm. **(C)** As an example, 10,000 single molecule points were simulated on an ideal radius (R_I_) of 25 nm and given a localization error based on the precision distribution from (E). Grayscale values indicate precision assigned to each single molecule. **(D)** The 3D transformation from (F) shows that the spread of each peak is approximately equal to the weighted average or “effective” precision (σ_EP_) of the distribution rather than the cutoff precision.

**Supplementary Figure 3.**
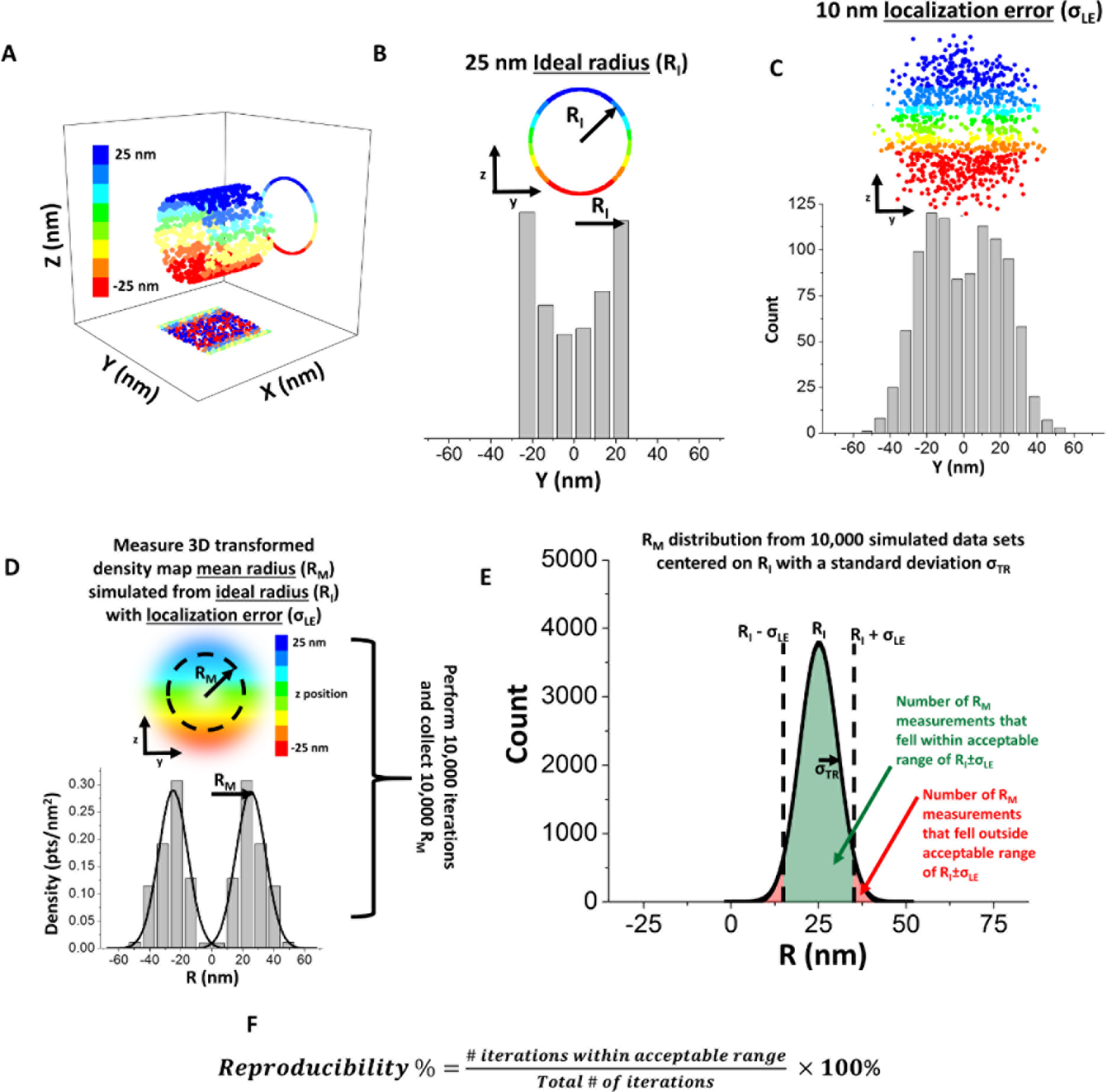
SPEED microscopy and 3D transformation reproducibility percentage using a simulation-based approach. For any given set of simulated data, the bin size is varied from 1 nm to the precision that is 10 nm in this example. **(A)** Data sets were simulated in three dimensions. Color bar indicates z position of the simulated points. **(B)** Each data set was simulated first with an ideal 25-nm radius (RI). (C) Subsequently, a localization error (σ_LE_) of 10 nm was added to each point. **(C)** Using a 5-nm bin size for demonstration, the 2D histogram of the simulated data set with a 25-nm radius and 10-nm localization precision was determined. **(D)** 10,000 data sets were simulated with an ideal 25-nm radius (R_I_) and a localization error (σ_LE_) of nm. The resultant 3D histograms were then each fitted with a Gaussian function to localize the mean position of each peak, which is designated as the mean radius R_M_. **(E)** The histogram for all the R_M_ values was determined and the number of simulated data sets that fell within the acceptable range of R_I_±σ_LE_ were counted. The acceptable range of R_I_±σ_LE_ was chosen because, in principle, the Rayleigh criterion limited the resolution of any single 3D histogram to the spread of that distribution, which was due to the simulated localization error (σ_LE_). After 10,000 simulations, the histogram for R_M_ values converges on the mean (R_I_) from which they were originally sampled, while the spread of the R_M_ histogram (σ_TR_) converges on a value that is due to the number of simulated points in each distribution and simulated localization error. **(F)** Reproducibility percentage was defined as the number of R_M_ values that fell within the acceptable range of R_I_±σ_LE_ divided by the total number of simulated data sets and multiplied by 100%.

**Supplementary Figure 4.**
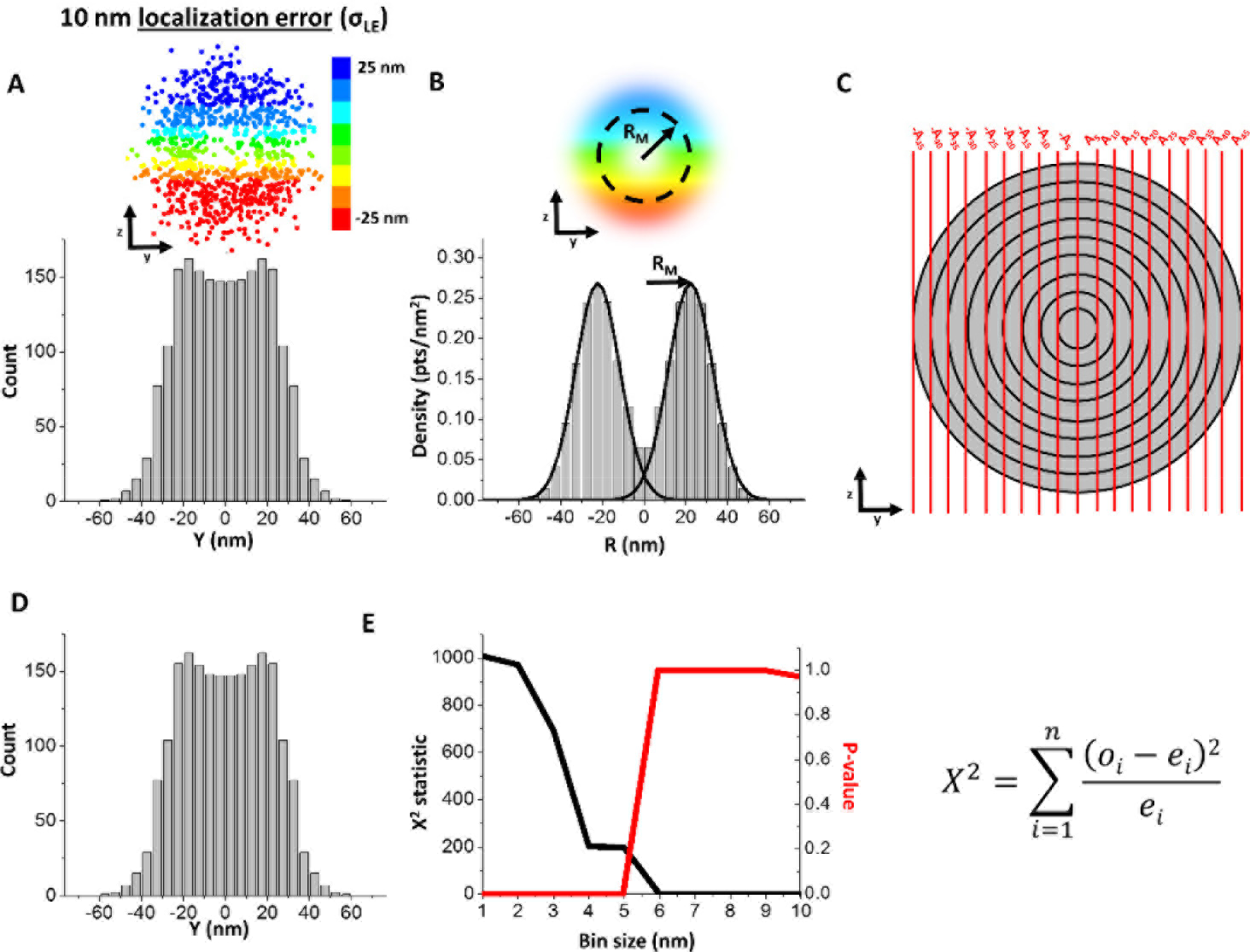
Optimal bin size determination using Chi-square error analysis. For any given set of simulated data, the bin size is varied from 1 nm to the precision that is 10 nm in this example. **(A)** Data sets were simulated in three dimensions. Color bar indicates z position of the simulated points. Each data set was simulated first with an ideal 25-nm radius (R_I_). Subsequently, a localization error (σ_LE_) of 10 nm was added to each point. Using a 5-nm bin size for demonstration, the 2D histogram of a simulated data set with a 25-nm radius and 10-nm localization precision was determined. **(B)** The 3D density histogram was then obtained via the 2D to 3D transformation algorithm and the peaks were fit with Gaussian distributions. **(C)** The 5nm bin size area matrix was calculated and multiplied by the 3D density distribution in (C) to reconstruct the 2D distribution. **(D)** The values of the reconstructed 2D distribution were then compared bin-by-bin to the original 2D distribution (as shown in A) using the Chi-square analysis equation where ‘o’ refers to the observed histogram values in (D), ‘e’ refers to the expected histogram values in (A), ‘i’ refers to the bin, and ‘n’ refers to the total number of bins with histogram values in them. **(E)** The Chi-square statistic and p-value were then plotted across the potential bin size values. A p-value ≤ 0.05 indicates that the 2D histograms in (A) and (D) are different from each other, suggesting the lack of enough data to allow sufficient sampling from each bin. A p-value > 0.05 indicates that the 2D histograms are not statistically different and likely have enough data points to accurately measure the value in each bin. Chi-square analysis was performed 10 times for each set of simulation parameters.

**Supplementary Figure 5.**
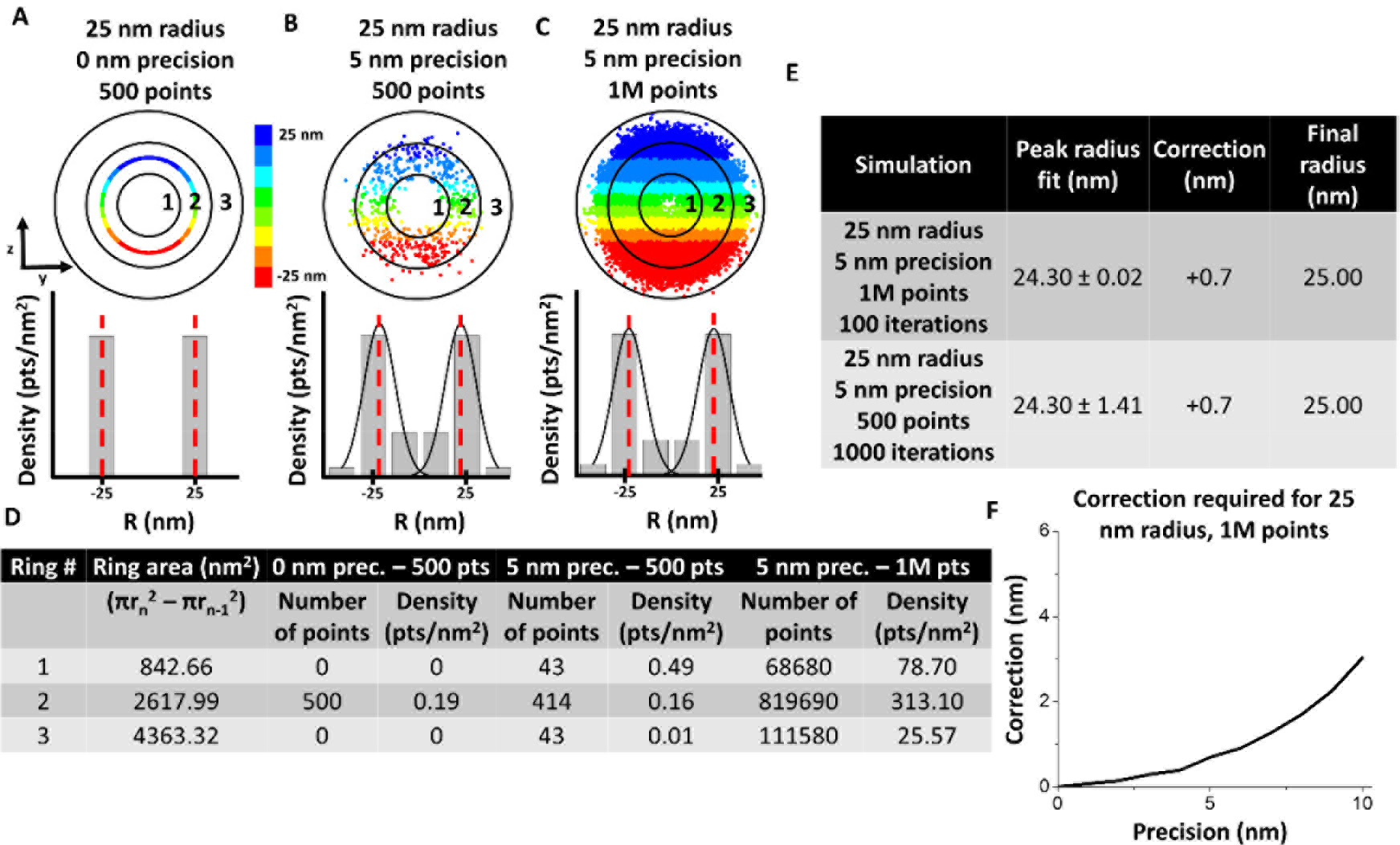
Sensitivity of inner bins necessitates slight correction of peak position during simulation. **(A), (B), and (C)** Simulated single molecule data and corresponding 3D density histogram for simulation with 25 nm radius, 0 nm localization error, and 500 points. Red dashed lines indicate mean peak fitting. **(B)** Simulated single molecule data and corresponding 3D density histogram for simulation with 25 nm radius, 5 nm localization error, and 500 points. Red dashed lines indicate mean peak fitting. **(C)** Simulated single molecule data and corresponding 3D density histogram for simulation with 25 nm radius, 5 nm localization error, and 1 million points. Red dashed lines indicate mean peak fitting. **(D)** Table showing the calculation to obtain each bin of the 3D density histogram. **(E)** Table showing that even one million points does reconstruct a precise 25 nm peak fitting due to the fact that the inner radial bins have smaller area and are slightly more sensitive to changes in density. **(F)** Correction required for each precision up to 10 nm for a 25 nm radius. This correction process was performed before each simulation to accurately localize the R_M_ density peak and correlate it to the ideal R_I_ from which the data was simulated.

**Supplementary Figure 6.**
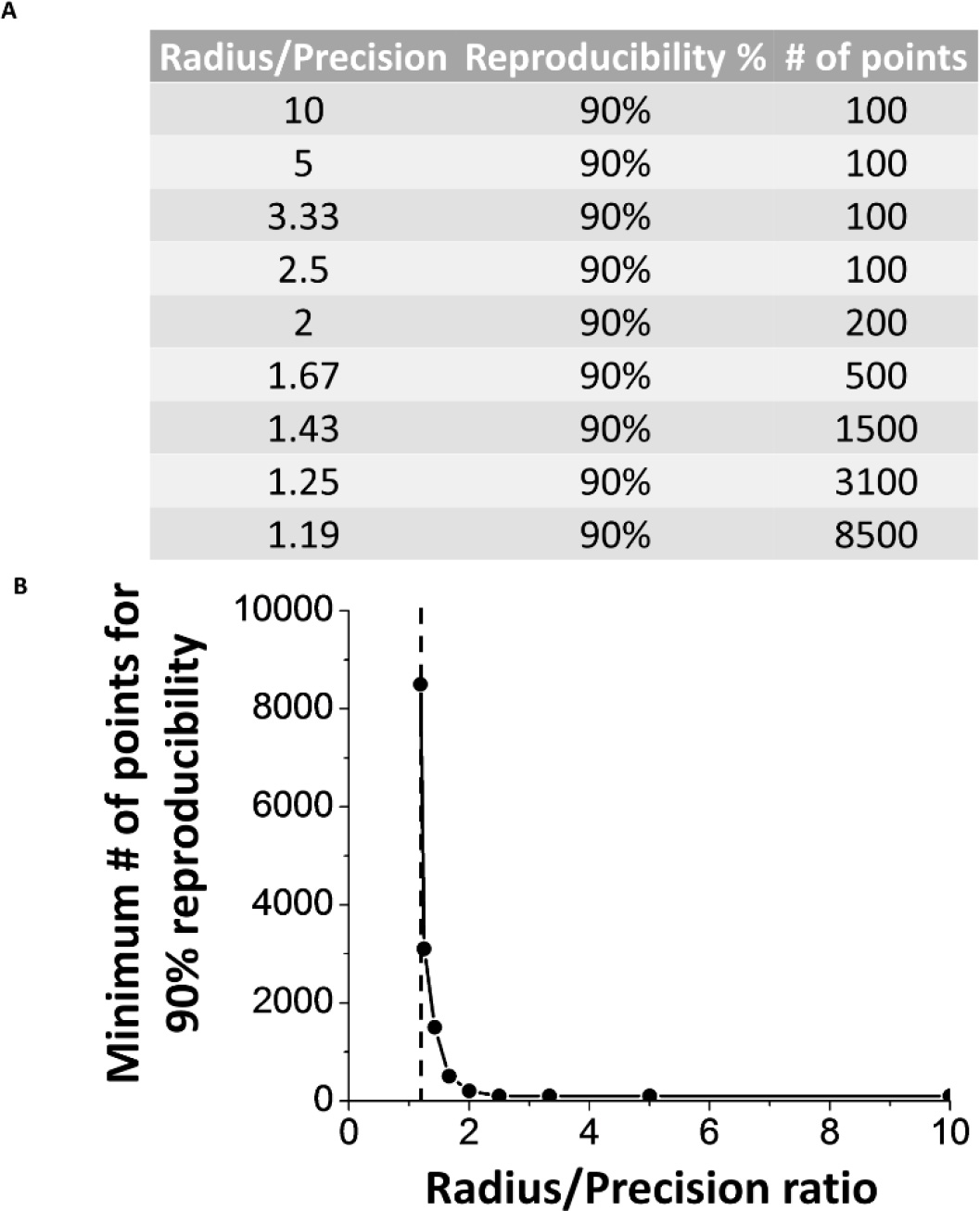
The minimum number of points required to resolve a transport route above the radius/precision threshold. **(A)** Table illustrating the minimum point number requirements for 90% reproducibility across a range of radius/s.d. ratios which covers the experimental data in Figure 4. Point numbers were rounded to the nearest 100 points. % route precision was calculated by dividing the transport route localization error (σ_TR_) by the ideal radius (R_I_). **(B)** Graph showing the table from (A) plotted in black dots and solid lines down to the minimum distinguishable R/P ratio threshold of 1.19 (dashed line). The green region of the graph represents the R/P ratio range that can distinguish a transport route with an empty center (bimodal distribution in 3D density map; Figure 2) from a transport route with a filled center (single central peak in 3D density map; Figure 2). Colored dots represent the location of R/Pratio from experimental data from Figure 4. In the case of importin β1 (R/P=2.3-3.65; 450 points), tubulin (R/P=5.09; 112 points), and SSTR3 (R/P=7.34-12.7; 260 points), excess points above the threshold (100 points) were collected to ensure accurate localization of the transport routes. In the case of the GNC (3000 points), single molecule localization precision ≤ 5 nm, an R/P ratio of 0, and the hollow structure of the GNC suggest that the accurate interpretation of the data are that it likely has a single central transport route.

## Acknowledgements

We acknowledge Drs. L.P. Zweifel and R.Y.H. Lim (University of Basel, Switzerland) for the glass nanocapillary experiments and Dr. Bo Huang (University of California – San Francisco, USA) for his critical reading of this manuscript. The project was supported by grants from the National Institutes of Health (NIH GM097037, GM116204 and RGM122552A to W.Y.).

### Author contributions

A.R., J.K., W.L., and W.Y. designed experiments; A.R., J.K., W.L., and W.Y. performed singlemolecule tracking and super-resolution SPEED microscopy experiments; A.R., J.K., and W.L. prepared plasmids and established cell lines; A.R., J.K., W.L., and W.Y. conducted data analysis; A.R., W.L. and W.Y. wrote the manuscript.

## References

1 Huang, B., Bates, M. & Zhuang, X. Super-resolution fluorescence microscopy. Annual review of biochemistry 78, 993–1016 (2009).

2 Leung, B. O. & Chou, K. C. Review of super-resolution fluorescence microscopy for biology. Applied spectroscopy 65, 967–980 (2011).

3 Hell, S. W. & Wichmann, J. Breaking the diffraction resolution limit by stimulated emission: stimulated-emission-depletion fluorescence microscopy. Optics letters 19, 780782– (1994).

4 Betzig, E. et al. Imaging intracellular fluorescent proteins at nanometer resolution. Science 313, 1642–1645 (2006).

5 Akey, C. W. Interactions and structure of the nuclear pore complex revealed by cryo-electron microscopy. The Journal of Cell Biology 109, 955–970 (1989).

6 Akey, C. W. & Radermacher, M. Architecture of the Xenopus nuclear pore complex revealed by three-dimensional cryo-electron microscopy. The Journal of Cell Biology 122, 1–19 (1993).

7 Yang, W. & Musser, S. M. Nuclear import time and transport efficiency depend on importin β concentration. The Journal of cell biology 174, 951–961 (2006).

8 Pavani, S. R. P. et al. Three-dimensional, single-molecule fluorescence imaging beyond the diffraction limit by using a double-helix point spread function. Proceedings of the National Academy of Sciences 106, 2995–2999 (2009).

9 Bohm, U., Hell, S. W. & Schmidt, R. 4Pi-RESOLFT nanoscopy. Nature communications 7 (2016).

10 Chen, F., Tillberg, P. W. & Boyden, E. S. Expansion microscopy. Science 347, 543–548 (2015).

11 Chang, J.-B. et al. Iterative expansion microscopy. Nature 201, 7– (2017).

12 Whelan, D. R. & Bell, T. D. Super-resolution single-molecule localization microscopy: tricks of the trade. The journal of physical chemistry letters 6, 374–382 (2015).

13 Saxton, M. J. in Fundamental concepts in biophysics 1–33 (Springer, 2009).

14 Ma, J., Goryaynov, A., Sarma, A. & Yang, W. Self-regulated viscous channel in the nuclear pore complex. Proceedings of the National Academy of Sciences 109, 7326–7331 (2012).

15 Ma, J. & Yang, W. Three-dimensional distribution of transient interactions in the nuclear pore complex obtained from single-molecule snapshots. Proceedings of the National Academy of Sciences 107, 7305–7310 (2010).

16 Marshall, W. F. & Nonaka, S. Cilia: tuning in to the cell's antenna. Current Biology 16, R604–R614 (2006).

17 Yang, T. T. et al. Superresolution pattern recognition reveals the architectural map of the ciliary transition zone. Scientific reports 5, 14096 (2015).

18 Scholey, J. M. & Anderson, K. V. Intraflagellar transport and cilium-based signaling. Cell 125, 439–442 (2006).

19 Mooney, C. Z. Monte carlo simulation. Vol. 116 (Sage Publications, 1997).

20 Mahadevan, S. Monte carlo simulation. MECHANICAL ENGINEERING-NEW YORK AND BASEL-MARCEL DEKKER-, 123–146 (1997).

21 Thompson, R. E., Larson, D. R. & Webb, W. W. Precise nanometer localization analysis for individual fluorescent probes. Biophysical journal 82, 2775–2783 (2002).

22 Thompson, M. A., Biteen, J. S., Lord, S. J., Conley, N. R. & Moerner, W. Chapter Two-Molecules and Methods for Super-Resolution Imaging. Methods in enzymology 475, 2759– (2010).

23 Ram, S., Ward, E. S. & Ober, R. J. Beyond Rayleigh's criterion: a resolution measure with application to single-molecule microscopy. Proceedings of the National Academy of Sciences of the United States of America 103, 4457–4462 (2006).

24 Axelrod, D. Total internal reflection fluorescence microscopy in cell biology. Traffic 2, 764–774 (2001).

25 Deschout, H., Neyts, K. & Braeckmans, K. The influence of movement on the localization precision of sub-resolution particles in fluorescence microscopy. Journal of biophotonics 5, 97–109 (2012).

26 Wright, A., Bubb, W. A., Hawkins, C. L. & Davies, M. J. Singlet Oxygen-mediated Protein Oxidation: Evidence for the Formation of Reactive Side Chain Peroxides on Tyrosine Residues¶. Photochemistry and photobiology 76, 35–46 (2002).

27 Bernas, T., ZarpBski, M., Cook, R. & Dobrucki, J. Minimizing photobleaching during confocal microscopy of fluorescent probes bound to chromatin: role of anoxia and photon flux. Journal of microscopy 215, 281–296 (2004).

28 Song, L., Varma, C., Verhoeven, J. & Tanke, H. J. Influence of the triplet excited state on the photobleaching kinetics of fluorescein in microscopy. Biophysical journal 70, 29592968– (1996).

29 Zweifel, L. P., Shorubalko, I. & Lim, R. Y. Helium scanning transmission ion microscopy and electrical characterization of glass nanocapillaries with reproducible tip geometries. ACS nano 10, 1918–1925 (2016).

30 Hlawacek, G., Veligura, V., van Gastel, R. & Poelsema, B. Helium ion microscopy. Journal of Vacuum Science & Technology B, Nanotechnology and Microelectronics: Materials, Processing, Measurement, and Phenomena 32, 020801– (2014).

31 Weis, K. Regulating access to the genome: nucleocytoplasmic transport throughout the cell cycle. Cell 112, 441–451 (2003).

32 Fried, H. & Kutay, U. Nucleocytoplasmic transport: taking an inventory. Cellular and molecular life sciences 60, 1659–1688 (2003).

33 Fahrenkrog, B. & Aebi, U. The nuclear pore complex: nucleocytoplasmic transport and beyond. Nature reviews. Molecular cell biology 4, 757– (2003).

34 Händel, M. et al. Selective targeting of somatostatin receptor 3 to neuronal cilia. Neuroscience 89, 909–926 (1999).

35 Sharma, K., Patel, Y. C. & Srikant, C. B. Subtype-selective induction of wild-type p53 and apoptosis, but not cell cycle arrest, by human somatostatin receptor 3. Molecular Endocrinology 10, 1688–1696 (1996).

36 Florio, T. et al. Somatostatin inhibits tumor angiogenesis and growth via somatostatin receptor-3-mediated regulation of endothelial nitric oxide synthase and mitogen-activated protein kinase activities. Endocrinology 144, 1574–1584 (2003).

37 Ye, F. et al. Single molecule imaging reveals a major role for diffusion in the exploration of ciliary space by signaling receptors. Elife 2, e00654 (2013).

38 Borisy, G. et al. Microtubules: 50 years on from the discovery of tubulin. Nature reviews. Molecular cell biology 17, 322– (2016).

39 Nogales, E. Structural insights into microtubule function. Annual review of biophysics and biomolecular structure 30, 397–420 (2001).

40 Desai, A. & Mitchison, T. J. Microtubule polymerization dynamics. Annual review of cell and developmental biology 13, 83–117 (1997).

41 Sage, D. et al. Quantitative evaluation of software packages for single-molecule localization microscopy. Nature methods 12, 717– (2015).

42 Weber, K., Rathke, P. C. & Osborn, M. Cytoplasmic microtubular images in glutaraldehyde-fixed tissue culture cells by electron microscopy and by immunofluorescence microscopy. Proceedings of the National Academy of Sciences 75, 1820–1824 (1978).

43 Mortensen, K. I., Churchman, L. S., Spudich, J. A. & Flyvbjerg, H. Optimized localization analysis for single-molecule tracking and super-resolution microscopy. Nature methods 7, 377–381 (2010).

44 Quan, T., Zeng, S. & Huang, Z.-L. Localization capability and limitation of electron-multiplying charge-coupled, scientific complementary metal-oxide semiconductor, and charge-coupled devices for superresolution imaging. Journal of biomedical optics 15, 066005–066005–066006 (2010).

45 Robbins, M. S. & Hadwen, B. J. The noise performance of electron multiplying charge-coupled devices. IEEE transactions on Electron Devices 50, 1227–1232 (2003).

46 von Appen, A. et al. In situ structural analysis of the human nuclear pore complex. Nature 526, 140– (2015).

47 Protein Data Bank Japan (PDBj). Date accessed: 8/3/2017

48 Blair, H. J. et al. Evc2 is a positive modulator of Hedgehog signalling that interacts with Evc at the cilia membrane and is also found in the nucleus. BMC biology 9, 14 (2011).

49 https://openi.nlm.nih.gov. Date accessed: 8/3/2017. This was modified from original version. License: https://creativecommons.org/licenses/by/2.0/

50 Ti, S.-C. et al. Mutations in human tubulin proximal to the kinesin-binding site alter dynamic instability at microtubule plus-and minus-ends. Developmental cell 37, 72–84 (2016).

51 Czarnecki, P. G. & Shah, J. V. The ciliary transition zone: from morphology and molecules to medicine. Trends in cell biology 22, 201–210 (2012).

